# Neural mass modelling for the masses: Democratising access to whole-brain biophysical modelling with FastDMF

**DOI:** 10.1101/2022.04.11.487903

**Authors:** Rubén Herzog, Pedro A.M. Mediano, Fernando E. Rosas, Andrea I. Luppi, Yonatan Sanz Perl, Enzo Tagliazucchi, Morten Kringelbach, Rodrigo Cofré, Gustavo Deco

## Abstract

Different whole-brain models constrained by neuroimaging data have been developed during the last years to investigate causal hypotheses related to brain mechanisms. Among these, the Dynamic Mean Field (DMF) model is a particularly attractive model, combining a biophysically realistic single-neuron model that is scaled up via a mean-field approach and multimodal imaging data. Despite these favourable features, an important barrier for a widespread usage of the DMF model is that current implementations are computationally expensive – to the extent that the model often becomes unfeasible when no high-performance computing infrastructure is available. Furthermore, even when such computing structure is available, current implementations can only support simulations on brain parcellations that consider less than 100 brain regions. To remove these barriers, here we introduce a user-friendly and computationally-efficient implementation of the DMF model, which we call *FastDMF*, with the goal of making biophysical whole-brain modelling accessible to neuroscientists worldwide. By leveraging a suit of analytical and numerical advances – including a novel estimation of the feedback inhibition control parameter, and a Bayesian optimisation algorithm – the FastDMF circumvents various computational bottlenecks of previous implementations. An evaluation of the performance of the FastDMF showed that it can attain a significantly faster performance than previous implementations while reducing the memory consumption by several orders of magnitude. Thanks to these computational advances, FastDMF makes it possible to increase the number of simulated regions by one order of magnitude: we found good agreement between empirical and simulated functional MRI data parcellated at two different spatial scales (N=90 and N=1000 brain regions). These advances open the way to the widespread use of biophysically grounded whole-brain models for understanding the interplay among anatomy, function and brain dynamics in health and disease, and to provide mechanistic explanations of recent results obtained from empirical fine-grained neuroimaging data sets, such as turbulence or connectome harmonics.

**Highlights:**

- We present the FastDMF, a user-friendly and computationally efficient implementation of the Dynamic Mean Field model for simulations of whole-brain dynamics.
- Using analytical and numerical tools, we develop a novel estimation of the feedback inhibition control based on the structural connectivity, bypassing an important computational bottleneck.
- The FastDMF is coupled with a Bayesian Optimization algorithm significantly reducing the number of simulations required to fit the FastDMF to empirical neuroimaging data.
- Our advances open the possibility of simulating thousands of brain regions in a biophysically grounded whole-brain model.

## Introduction

Recent advances in non-invasive brain-imaging technology provide a fertile ground to investigate how the anatomical structure of the brain shapes complex neural dynamics, in both healthy and pathological conditions. As a matter of fact, the high versatility of the available imaging modalities has triggered a plethora of research efforts, which in turn have delivered important advances in human neuroscience^1–3^.

To move beyond correlational inference and towards causal understanding, such tools can be combined with causal manipulations of brain activity induced by tasks, pharmacology, or non-invasive brain stimulation (e.g., TMS, tDCS), and in rare cases direct cortical stimulation in surgical patients^4^ or the study of other neuropsychiatric patients^5–8^. However, ethical and practical considerations mean that such approaches fall short of the exquisitely fine-grained experimental control that is nowadays available in animal models, which has provided fundamental new insights about the causal mechanisms of brain function^9^.

One attractive approach to bridge the gap between neuroimaging data and causal mechanisms in humans is whole-brain modeling^10–17^, where neurobiologically-inspired models informed by multimodal empirical data are used to reverse-engineer various aspects of brain function. Whole-brain models typically consist of equations describing local dynamics of neuronal populations, coupled through a network of empirically-derived anatomical connections (e.g., tract-tracing data from animals, or diffusion tensor imaging (DTI) data from humans)^10–12,18–21^. After appropriate tuning, such models make it possible to simulate plausible brain dynamics, which can then be compared with empirical brain dynamics (as measured e.g. with functional MRI or EEG), thereby allowing researchers to investigate the relationship between anatomical connectivity and brain activity and local network dynamics, providing a valuable tool to bridge scales and narrow down the space of mechanistic explanations compatible with empirical findings.^3,22^ Whole-brain models also provide an ethical and inexpensive “digital scalpel” that allows researchers to explore the counterfactual consequences of modifying structural or dynamical aspects of the brain, some of which would be hard — if not entirely impossible — to assess experimentally. This ability to assess counterfactuals makes whole-brain modeling a promising tool to deepen our understanding of the mechanisms behind brain disorders, and to explore novel therapeutic interventions, including drug treatments or brain stimulation^23^.

The literature offers a wide range of whole-brain models, with some of them being supported on platforms such as *The Virtual Brain*^24^ or *neurolib*^25^. Available models can be classified according to their trade-off between conceptual simplicity and neurobiological realism. On one side of this spectrum, models prioritize local equations that reproduce the empirical data with a minimal number of assumptions and fine-tuning of parameters, and thus are useful to reveal general dynamical principles (e.g. simulating brain areas as nonlinear oscillators near the transition to global synchronization). On the other side of the spectrum, models are more complex but they can be interpreted in terms of the biology and biophysics of neurons (e.g. modeling brain areas as balanced excitatory/inhibitory neural populations)^10^. In general, the former are faster to simulate and easier to tune and control, while the latter provide more realistic explanations and can incorporate additional neurobiological information, such as receptor densities obtained using positron emission tomography (PET) or transcriptomic data.^26^

One of the most prominent biophysically-grounded whole-brain models is the Dynamic Mean-Field (DMF) model^10,27^, which results from applying a mean-field approach to single neuron models, allowing to model the dynamics of the mean firing rate of a macroscopic brain region^28,29^. This approach allows to model the local dynamics of each brain region as two interacting neural populations: one excitatory (E) and another inhibitory (I) (see Fig. 1B). In addition to interacting with its coupled I pool, the E pool receives/sends excitatory inputs/outputs from/to the E pools of other regions according to the connectome blueprint, which determines the weight of this inter-regional interaction (Fig. 1A). Finally, the simulated firing activity of each E pool is transformed into BOLD-like signals using the Balloon-Windkessel (BK) model^30^ (Fig. 1C), allowing a direct comparison with empirical fMRI data. The DMF has been used to simulate BOLD signals measured during an ample range of different conditions and global brain states^12,15,17,31–34^.

**Figure 1.**
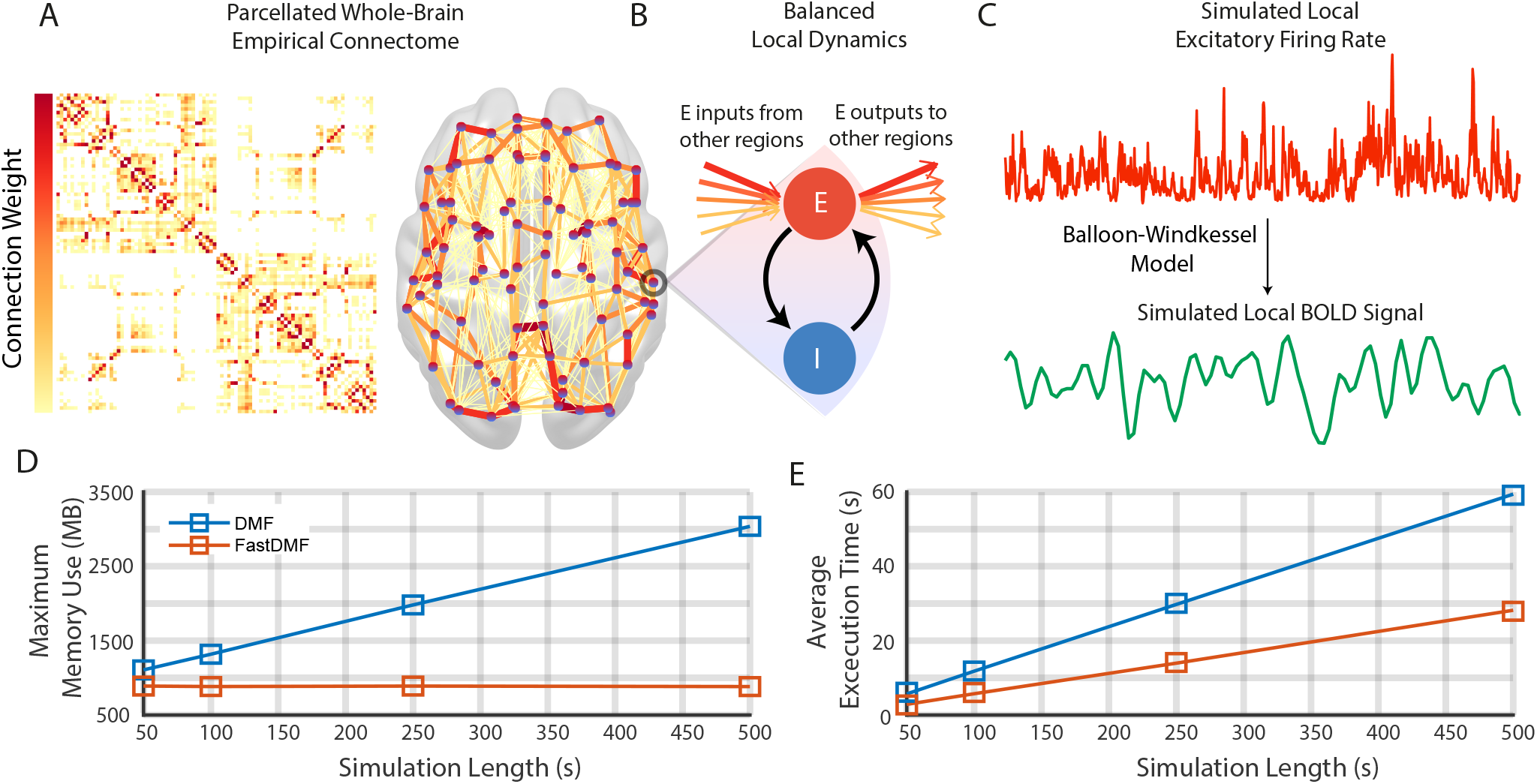
Simulations using the Fast Dynamic Mean Field Model. **(A)** Brain regions are defined by a parcellation (here, the AAL90) and their connections are empirically obtained from DTI (see Methods for details). In the schematic brain, circles are brain regions and connections between brain regions are colored according to their connection weight in the connectome. **(B)** The local dynamics of each brain region are simulated by recurrently connected pools of excitatory (E, red circle) and inhibitory (I, blue circle) neuronal populations. In turn, brain regions are connected through the E pool, such that the excitatory inputs from other regions are weighted according to their respective connection weight, and their sum is scaled by the Global Coupling parameter *G*. The connection from the I pool to the E one - *J*_*n*_, the local inhibitory feedback - compensates for the excess of excitatory activity injected from other regions. **(C)** The simulated firing rate of each excitatory pool is used to generate BOLD-like signals for each brain region using the Balloon-Windkessel model. **(D)** Maximum memory used by the current implementation (DMF) and by ours (FastDMF) to simulate BOLD signals of different length. Both implementations used the connectome shown in **A. (E)** Same than **D**, but for the average execution time among 10 repetitions.

Despite these achievements, a few limitations prevent the DMF model from being more widely exploited by the scientific community. The main limitation is that current publicly available implementations of the DMF model are computationally expensive, with demanding time and memory requirements. This problem is exacerbated by two further aspects of the DMF: 1) the need to calibrate the model’s Feedback Inhibition Control (FIC) parameters, which stabilise the firing rates of E pools, and have been shown to provide more realistic activity^10^ and richer dynamics^35^; and 2) the need to optimise the model’s free parameters to fit statistical features of a given empirical dataset. Both aspects involve running a large number of long simulations and thus inflate the overall computational costs, which often make the requirements of the DMF prohibitive for researchers with no access to high-performance computing infrastructure. This represents an undue obstacle to the ability of neuroscientists across the globe to contribute to brain modelling research: the choice of an appropriate computational model should depend only on its suitability to address specific research question, rather than being contingent on the availability of high-performance resources.

Moreover, the high computational burden of the current implementations of the DMF does not only restrict its usage, but also severely restricts the spatial resolution of what can be simulated. The existent implementations of the DMF are based on parcellations with less than 100 regions, –typically given by the Automatic Anatomical Labelling (AAL90)^36^ or the Desikan-Killiany^37^ parcellation– because, otherwise, it would be unfeasible to simulate brain activity based on fine-grained parcellations. However, recent advances in neuroimaging data analysis involve measuring brain activity at multiple spatial scales, from coarse to fine-grained, revealing new insights about underlying brain dynamics^38^ and its relevant operational scales^39^. Therefore, being restricted to simulating biophysically realistic brain activity at a coarse-grained spatial scale represents another barrier that hinders the ability of neuroscientists to leverage the full potential of DMF modelling.

To address both of these limitations, here we present *FastDMF*: a time- and memory-efficient implementation of the DMF model, which reduces its computational demands so it can be run and fit efficiently on any contemporary desktop computer, dramatically widening access to this kind of computational modelling approach. The FastDMF implementation is built by leveraging a number of key advances:

- It provides an improved implementation of the DMF model, which is significantly faster and reduces memory consumption by several orders of magnitude.
- It uses a novel connectome-dependent local inhibitory feedback mechanism, which replaces the standard FIC optimization problem and radically reduces the number of FIC calibration parameters.
- It leverages a Bayesian optimization algorithm, which substantially reduces the number of simulations required to fit the model to empirical data. This algorithm performs a smart sampling of the parameter space instead of the grid search used in previous approaches^12,33^.

These advances make FastDMF a computationally efficient, easily accessible, and biophysically grounded whole-brain model.

To showcase the advantages of the FastDMF, we first compare our implementation to the current one (published with Ref^12^) in terms of computational costs. Then, we show analytical and numerical evidence supporting a connectome-dependent linear solution to the FIC optimization problem. Finally, using the Bayesian Optimization algorithm, we fit an fMRI dataset parcellated at two different spatial scales (N=90 and N=1000 regions). Using less than 350 optimization iterations, which can be performed in personal computers in matter of minutes, we achieve an excellent agreement between empirical and simulated data for both scales, showing the performance and broad applicability of the FastDMF to different scenarios. Furthermore, to facilitate the use of FastDMF by the community, we provide all codes that implement FastDMF. FastDMF is coded in C++ and is usable on both Matlab and Python and can be found on https://gitlab.com/concog/fastdmf.

## Results

### Fast and efficient computational implementation

The first advancement to make the FastDMF widely usable is to provide an open-source implementation with two main advantages: faster execution time and less memory usage. This is achieved through three main improvements: i) The core of FastDMF is written in C++ and takes advantage of the advanced linear algebra library Eigen,^40^ which provides a fast and cross-platform toolkit for numerical operations. ii) The FastDMF and the Balloon-Windkessel hemodynamic (BK) function to compute the BOLD signals run in parallel, further reducing the overall execution time of the simulation. iii) The DMF and the Balloon-Windkessel numerical integrators access a shared memory via a simplified producer-consumer architecture, which, given the large difference in timescale between the BOLD and the firing rate signals, allows FastDMF to radically reduce memory usage through shared finite-size buffer.^1^

To benchmark our implementation, we ran simulations of varying lengths using the AAL90 structural connectome^36^ and measured both memory usage and execution time. As expected, FastDMF was faster and more memory-efficient than the public Matlab implementation^12^. Thanks to the circular buffer, memory is effectively constant for all simulation times (Fig. 1D) and runtime per second of simulated activity is less than half of the Matlab DMF (Fig. 1E).

Finally, in addition to the underlying C++ implementation, the FastDMF library incorporates interfaces for both Matlab and Python, using the Mex^41^ and Boost Python^42^ libraries respectively. These enable researchers to run and analyse simulations from a language of their choice, thus making the integration of FastDMF in their existing pipelines easier.

### Linear solution to the FIC optimization problem

The second main improvement was to provide a linear solution to the optimization of the Feedback Inhibition Control^10^ (FIC). In brief, the Global Coupling (*G*) parameter of the DMF scales the inputs that each region receives from the rest of the network, allowing to tune the model closer/farther to an optimal working point, where some desired statistical feature of the empirical neuroimaging data is reproduced, such as the Functional Connectivity (FC) or the Functional Connectivity Dynamics (FCD). To compensate for the excitation that each pool receives from the other regions via the connectome, a FIC parameter is optimized through recursive adjustments to clamp the firing rate within a neurobiologically plausible range of 3-4 Hz for each local excitatory neural population, preventing the system from entering a hyper-excitation regime. The solution to this problem, based in an iterative process of increasing and decreasing the local inhibition until convergence, is explained in detail in Ref.^10^, and it is implemented in the scripts accompanying the publication of Ref.^12^ and gives the optimal local inhibitory feedback *N* − dimensional vector (*J*^opt^) for a given *G*, where *N* is the number of regions. However, for a standard setup these scripts took more than 2 months on a 32-core cluster to converge. Even worse, running the model using different connectivity matrices would require to separately optimize the FIC for each one of them, severely hindering the endeavour of investigating how structure shapes function through biophysical DMF modelling.

Despite the lack of prior information regarding the expected *J*^opt^ in the DMF models, results from other type of whole-brain models suggest that *J*^opt^ is correlated with local connectivity measures of the structural connectivity^35^. Accordingly, here we show that a first-order analytical solution for a stationary state where the expected values of firing rates are fixed to 3.4 Hz (average excitatory firing rate for *G*=0) predicts that *J*^opt^ is region-specific, and its magnitude is proportional to *G* and to the local connectivity strength *β*_*n*_ := ∑_*p*_ *C*_*np*_ is the strength of node *n* computed from the anatomical connectivity matrix *C*.^2^ (see Methods).

We write this solution as

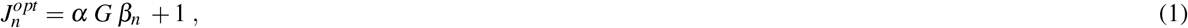

where *α* is a scaling factor that represents the global excitation-to-inhibition (E/I) ratio (see Methods). According to this approximation, for a fixed *G, α* is the only parameter to estimate in order to solve the feedback inhibitory control problem.

As a previous well-known and publicly available implementation of the DMF model Ref.^12^ used the AAL90 parcellation, we numerically tested our first-order approximation using this parcellation. After the optimization converged, for each *G*, we plotted 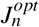 vs the local connectivity strength (*β*_*n*_), finding a linear relationship whose slope linearly increased with *G* (Fig. 2A,B). Then, to estimate *α* of Eq. 1, we found the scaling factor between the slope and *G*, with the constraint that the slope for *G* = 0 should be 0. We used least squares to find the optimal value of *α* (0.725). However, this value gives lower goodness of fit for higher *G* values, which are usually the values where the model better fits empirical data^10,12^. Accordingly, we used weighted least squares, giving 10 times more weight to *G* values larger than 2.1 (close to bifurcation with *α* = 0.725), finding an optimal value for *α* = 0.75. This approach, as expected, better matches the slope values for high *G* values and also extends the range where stability can be attained by the linear approximation (Fig. 2C, green dots).

**Figure 2.**
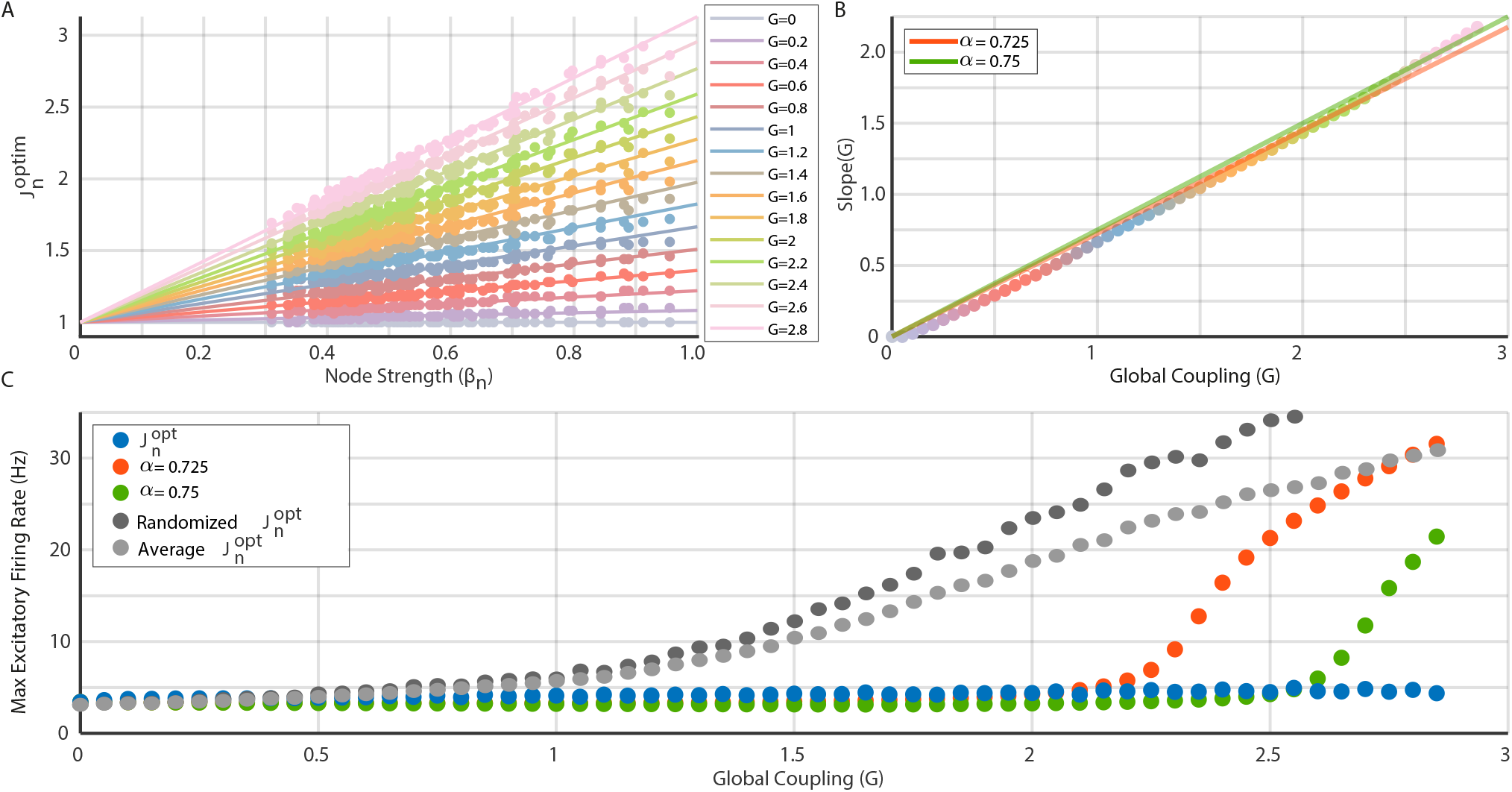
Optimal local inhibitory feedback depends on Global Coupling and local anatomical connectivity strength. **(A)** Optimal local inhibitory feedback 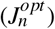 is well approximated by a linear function of region strength (*β*_*n*_) for a wide range of *G* values (colors). **(B)** The slope of the 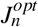 vs *β*_*n*_ scales linearly with the Global Coupling, giving an optimal slope of 0.725(red solid line), however, using 0.75 (green solid line) better matches the high *G* values. **(C)** 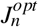 attains stability for all *G* values between [0,3], while the linear approximation using *α* = 0.725 diverges to the high excitability regime close to G=2 (red dots). Using *α* = 0.75 (green dots) attains stability in a larger range of *G* values, diverging at *G* ∼ 2.5. To illustrate the topographical specificity of 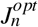, we used a randomized version of 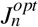 (dark gray circles) and an homogeneous *J*_*n*_ equal to the average 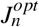 (light gray circles), both fails to control the global dynamics for increasing values of *G*.

Note that a second-order approximation can be also used for the relationship between 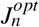, *G*, and *β*_*n*_, but our numerical simulations show that the range where stability is attained by second-order approximations is reduced, in comparison to the first-order one.

Finally, as a control, we used two modified versions of *J*^*opt*^ to run the DMF model: i) a randomized (shuffled) version of *J*^*opt*^, and ii) a homogeneous version, where *J*_*n*_ for all regions is equal to the average of *J*^*opt*^ (Fig. 2C). We found that both alternative versions of *J*^*opt*^ fail to attain stability for high values of *G >* 0.5), showing that *J*^*opt*^ is region-specific.

Our results alleviate the huge amount of simulations required to properly tune the FIC, reducing the number of free parameters of the optimization problem from N to one. In addition, we provide a biophysical interpretation of FIC in terms of the local connectivity strength and the global E/I balance.

### Reproducing the FCD of resting state fMRI data for two different spatial resolutions

The third major improvement was to implement a Bayesian Optimization algorithm to find the optimal set of parameters that reproduces a statistical feature of the empirical data (or more than one). The current DMF implementation finds the optimal working point by using a sub-optimal strategy based on grid search. Bayesian Optimization algorithms, on the other hand, estimate the objective function by sampling the parameter space efficiently, finding the minimum of a wide variety of functions^43,44^. Here, following previous applications of the DMF to fMRI data^12^, we chose as objective function the Kolmogorov-Smirnov (K-S) distance between the pooled empirical FCD distribution (*FCD*_*emp*_, computed using the FCD from all subjects) and the FCD histogram obtained with the DMF (*FCD*_*dm f*_).^45^ In addition, instead of first optimising FIC to clamp the firing rates, and then optimising *G* to reproduce the FCD, we jointly optimised *G* and *α*, expecting that the optimal working point also satisfies the firing rates constraints. We ran the optimization until convergence to K-S distances was comparable to previous studies^12,33^.

To exhibit the advantages of the FastDMF, in addition to the AAL90 parcellation used up to this point for consistency with previous work, we also obtained a different estimate of the healthy human connectome, based on high-resolution diffusion data^46^ parcellated according to most fine-grained scale of the Schaefer atlas, which comprises 1000 functionally defined cortical regions^47^ (see Methods). We also used the same parcellation to obtain the functional MRI timeseries, obtaining data sets in two different spatial scales.

We emphasise that the two connectomes employed for the following analyses were obtained from different groups of healthy individuals, with different acquisition parameters, different reconstruction methods and softwares, different atlases (anatomically *versus* functionally defined), different resolution (differing by one order of magnitude) and only one of which includes subcortical regions (AAL90). In other words, we chose to use two healthy connectomes that differ on virtually every relevant methodological dimension, to establish the general validity of our results.

Indeed, we found comparable values of goodness-of-fit for the two spatial scales (K-S ∼ 0.15, (Fig. 3A,F)). Note that a difference in the optimal *G* value between the two spatial scales is to be expected: not just because two different connectomes were used, but also mathematically because of the larger number of regions in the Schaefer1000, which in consequence increases the average *β*_*n*_, reducing the magnitude of the Global Coupling required to drive the system to a high-excitability regimen.

**Figure 3.**
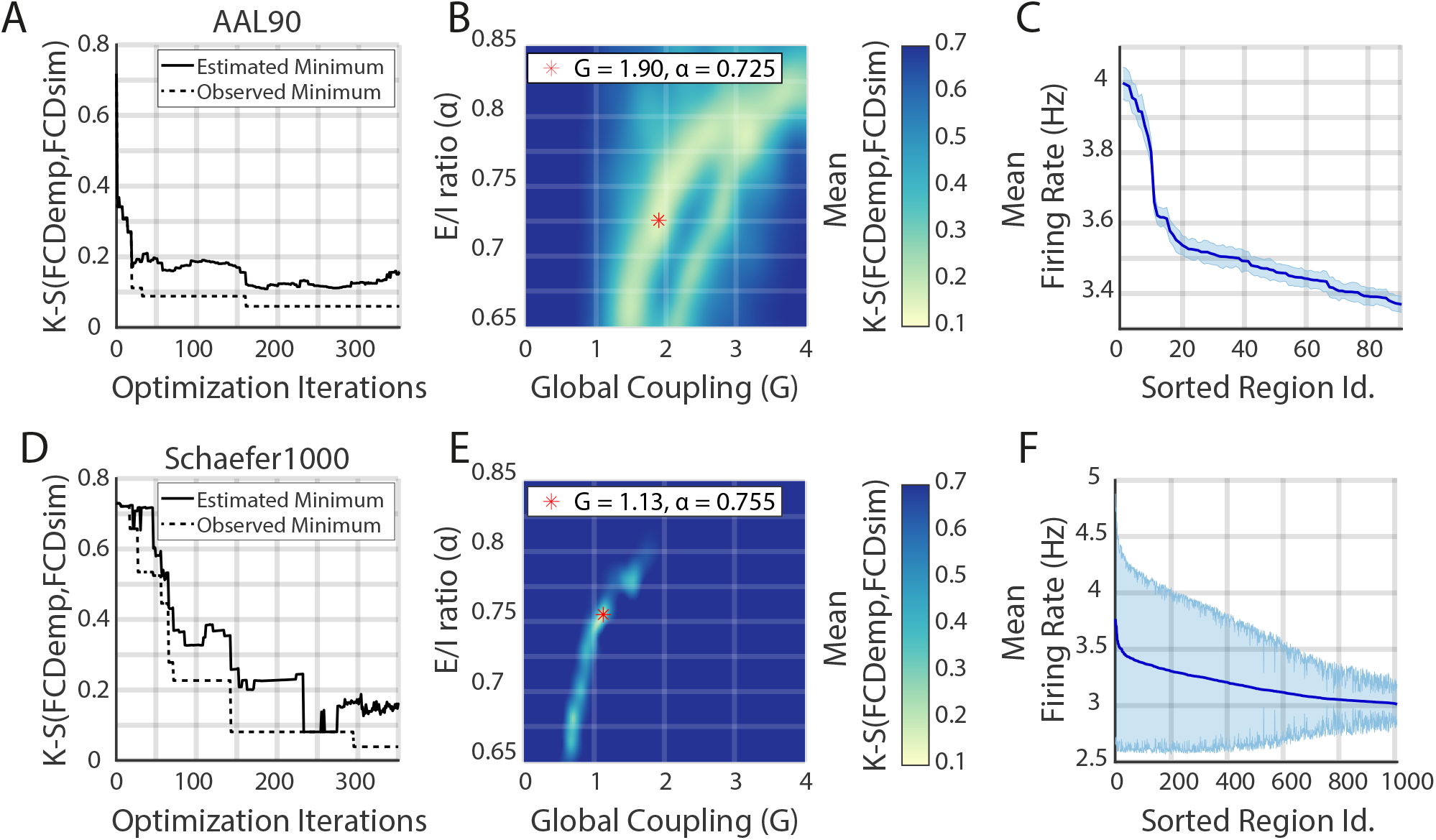
Fitting the FastDMF model to empirical data with Bayesian Optimization. **(A**,**D)** Convergence of the Bayesian optimization estimates of the minimum K-S distance (solid line), for the AAL90 and the Schaefer1000 parcellation, respectively. Dashed lines are the minimum observed K-S during the optimization process. **(B**,**E)** Bayesian optimization estimate of the mean K-S distance between the pooled empirical FCD (*FCD*_*emp*_) and the simulated FCD (*FCD*_*sim*_) as function of *G* and *α*. Red asterisk shows the optimal parameters. **(C**,**F)** Simulated brain regions sorted according to their average excitatory firing rate (solid line) with their respective standard deviation computed from 96 simulations (shaded area). All regions are within the expected range of firing rates.

Crucially, for both connectomes *α* is within the values shown in Fig. 2, demonstrating the robustness and generalisability of our estimation of this parameter, even when the empirical healthy connectomes used for the simulation originate from different cohorts and were obtained through different reconstruction approaches at different resolutions.

For both parcellations, we required 350 iterations to converge. Note that without this approach we would have to simulate a subset of parameter combinations several times, taking the risk of using a too coarse binning of the parameters space (to save time), thus missing the minimum. In addition, the Bayesian optimization algorithm can take advantage of parallel computing, making the optimization procedure even faster.

We further evaluated the Bayesian Optimization results by using the optimal parameters to run 96 simulations with different initial conditions and a simulation length comparable to resting state recordings (∼ 10 mins). First, we checked the average excitatory firing rates, finding that for both spatial scales they remained in the desired range (3-4 Hz, Fig. 3B,G)). However, given the increased number of regions of the Schaefer1000 parcellation, the average firing rates showed increased variance, reflecting more susceptibility to the initial conditions. Second, we checked that the best simulation closely matched the empirical FCD distribution (Fig. 4A,F), reproducing the bimodality of the empirical FCD distribution. Note that the resting-state Functional Connectivity matrix (FC) is also well captured (Fig. 4B,G), even when it was not included as an objective function in the fitting procedure. Finally, we computed the K-S between the FCD of each subject 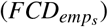 and the pooled FCD (*FCD*_*emp*_) to evaluate the goodness of fit respect to the variability of the empirical population. Accordingly, we computed the FCD between each simulation 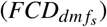 and *FCD*_*emp*_, finding a close match between both distributions for both spatial scales. As a final check, we also computed the mean squared error of the FC (MSE FC) as a measure of FC matrix similarity (see Methods). This results shows that the optimal parameters generate simulations whose K-S values and FC matrices are close to those obtained from experimental subjects (Fig. 4C,D,H,I).

**Figure 4.**
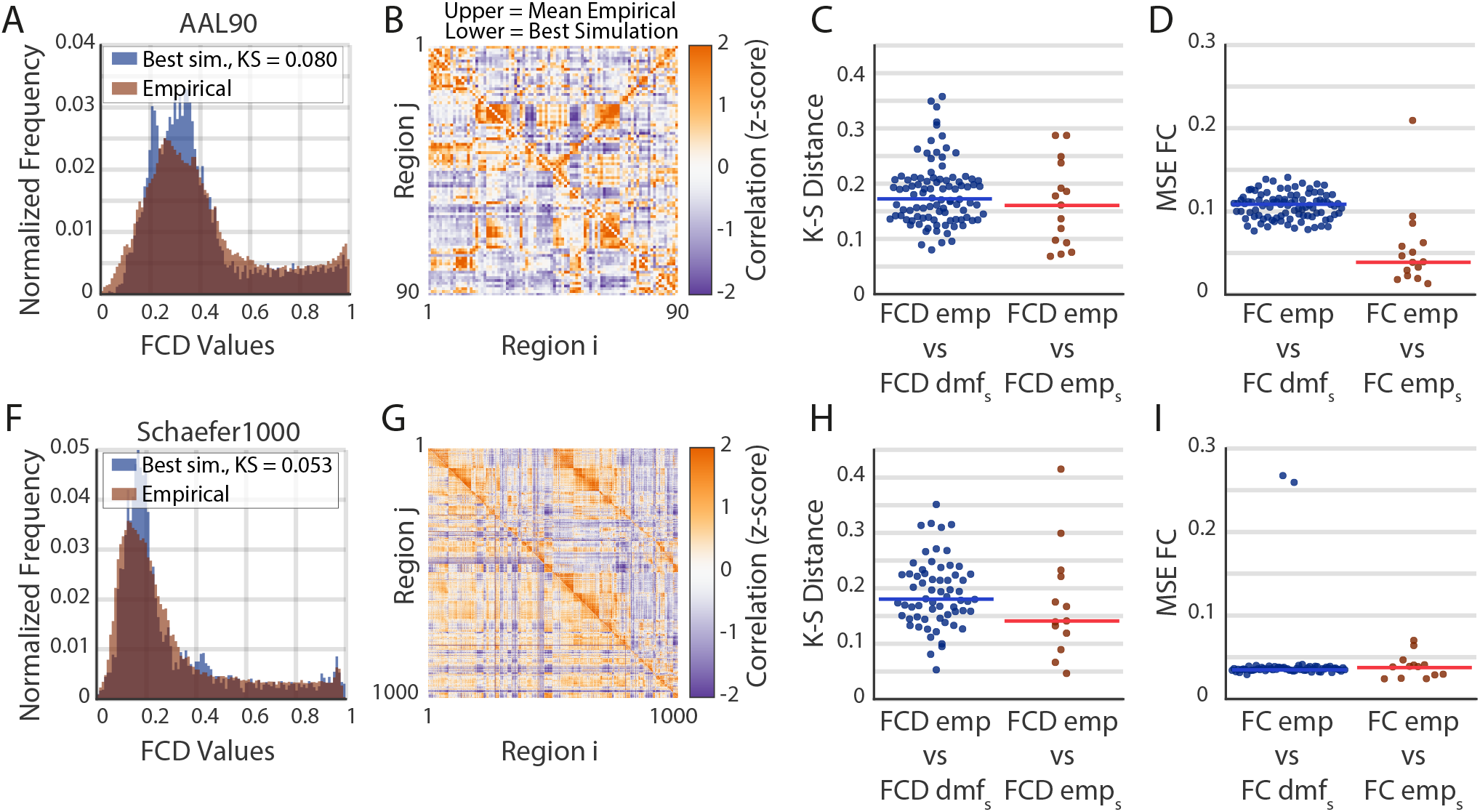
Comparison between empirical and simulated FCD and FC for two spatial scales. **(A**,**F)** FCD distribution for the pooled empirical data (red) and for the best simulation (blue) for the AAL90 and the Schaefer1000 parcellations, respectively. **(B**,**G)** Functional connectivity (FC) matrix for the average empirical (upper triangular) and for the best simulation (lower triangular). **(C**,**H)** K-S distance between simulated, 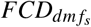, and pooled empirical *FCD*_*emp*_ (blue). The K-S distance between single subjects, 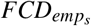, and pooled empirical *FCD*_*emp*_, is shown in red. Horizontal line is the median. **(D**,**I)** Same than (**C**,**H**), but for mean squared error of the FC matrix. In this case, *FC*_*emp*_ is the average FC between all subjects.

We highlight that this approach represents a considerable reduction in the number of simulations required to fit the model to empirical data, even more for the Schaefer1000 parcellation, which is prohibitive in current implementations. In addition, we remark that the optimal set of parameters that reproduces the empirical FCD also satisfies the local stability condition (i.e. stable firing rates), and also captures important structures of the FC matrix, showing the degree of generalization of the FastDMF model.

## Discussion

We introduced the FastDMF, an accessible, efficient, and biophysically grounded whole-brain model. Thanks to our combination of optimised implementation, efficient parameter search, and a novel method of estimating the feedback inhibitory control, we were able to fit empirical fMRI data using two parcellations representing drastically different spatial scales: the AAL90^36^ and the Schaefer1000^47^ parcellations, highlighting the flexibility of our approach and its robustness to a variety of methodological choices. These examples show that our method provides accurate results for a wide range of *G* parameter values, thus outperforming existing tools for computing FIC. In terms of applications, the FastDMF opens the way for biophysically grounded whole-brain models of a range of novel applications such as turbulence analysis, connectome harmonics, or detailed perturbation protocols, which requires detailed and coarse-grained descriptions. This application could lead to promising novel insights about brain dynamics and evoked responses.^38,48–51^.

Different alternatives have been proposed to model the FIC. A plasticity rule for a time-varying FIC has been used in a modified version of the DMF^52^, avoiding the optimization process, at the expense of increased model complexity. In fact, it has been suggested that the optimal FIC obtained by the iterative process Ref.^10^ corresponds to the integrated form of the time-varying FIC^35^. While our main purpose was to show the use of the FastDMF rather than explaining the possible mechanisms involved in the FIC or their potential functional implications, our model provides novel evidence in favor of a connection between the strength of the nodes in the connectome and FIC.

There is room for further improvement of our model. For example, a time-varying FIC could be implemented, following the approach proposed by Hellyer et al^35^. More general parameters allowing the incorporation of multiple receptor maps allowing to neuromodulate the activity of each brain region, principal gradient of gene expression or anatomical measures could be implemented in future developments.^53^ Additionally, new data modalities that can be derived from firing rates such as local field potentials or EEG-like could be future avenues of development.

The study of whole brain activity is an exciting research avenue that needs to be better developed to understand the causal interplay between brain structure and function in different brain health conditions, addressing the specific relevance of factors such as neuronal dynamics, neurotransmitter receptor density, and anatomical connectivity, among others. This, in turn, might accelerate brain research, identifying the biophysical mechanistic principles that relate different levels of brain organization, opening in this way the road for the development of new treatments to prevent and cure brain disease^3,23^. In line with this perspective, the FastDMF provides a general and accessible tool of simulation and analysis to be applied in multiple neuroimaging scenarios, that can be called from Python and Matlab, which should favor its use by the scientific community.

## Methods

### Dynamic mean-field model

The Dynamic Mean-Field (DMF) model introduced by Deco *et al*.,^10,12^ uses a set of coupled stochastic differential equations to model the dynamics of the activity at one or more interacting brain regions. In this model, each brain region is modelled as two coupled neuronal masses — one excitatory and one inhibitory — and considers excitatory and inhibitory synaptic currents mediated by NMDA and GABA_A_ receptors, respectively. Different brain regions (usually defined by a given brain parcellation) are coupled via their excitatory populations exclusively, according to the structural connectivity matrix.

The key idea behind the mean-field approximation is to reduce a high-dimensional system of randomly interacting elements to a system of elements treated as independent. This approach represents the average activity of a homogeneous population of neurons by the activity of a single unit of this class.

The model establishes that the *n*-th brain area obeys the following equations:

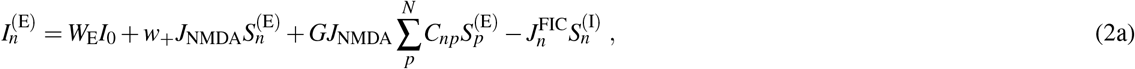

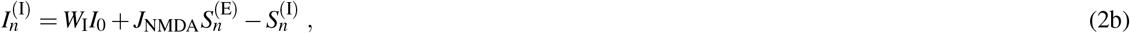

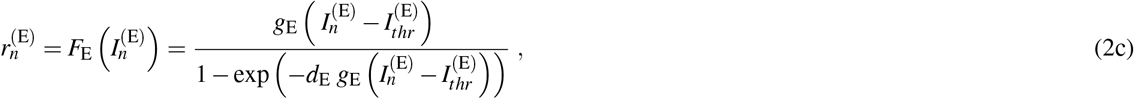

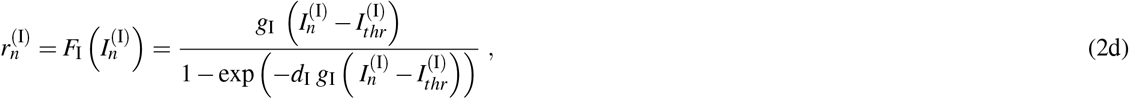

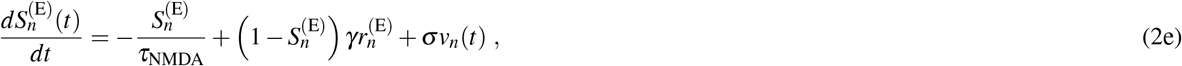

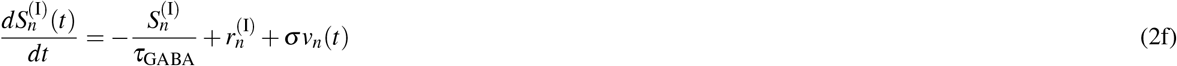

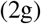

Above, for each excitatory (E) and inhibitory (I) neural mass, the quantities *I*_*n*_, *r*_*n*_, and *S*_*n*_ represent its total input current, firing rate, and synaptic gating variable, respectively. The functions *F*_E_(·), *F*_I_(·) determine the transfer functions (characterised by a *F-I curve*), representing the non-linear relationship between the input current and the output firing rate of excitatory and inhibitory neural populations. Crucial for our derivations, 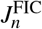 is the local feedback inhibitory control of region *n*, and *v*_*n*_ is uncorrelated Gaussian noise injected to region *n*^54^. The remaining quantities involved in these equations are introduced in Table 1. The interested reader is referred to the original publication for further details.^10^

**Table 1.** Dynamic Mean Field (DMF) model parameters

| *Parameter* | *Symbol* | *Value* |
| --- | --- | --- |
| External current | $I_0$ | 0.382 nA |
| Excitatory scaling factor for $I_0$ | $W_E$ | 1 |
| Inhibitory scaling factor for $I_0$ | $W_I$ | 0.7 |
| Local excitatory recurrence | $w_+$ | 1.4 |
| Excitatory synaptic coupling | $J_{NMDA}$ | 0.15 nA |
| Threshold for $F(I_n^{(E)})$ | $I_{thr}^{(E)}$ | 0.403 nA |
| Threshold for $F(I_n^{(I)})$ | $I_{thr}^{(I)}$ | 0.288 nA |
| Gain factor of $F(I_n^{(E)})$ | $g_E$ | 310 nC<sup>-1</sup> |
| Gain factor of $F(I_n^{(I)})$ | $g_I$ | 615 nC<sup>-1</sup> |
| Shape of $F(I_n^{(E)})$ around $I_{thr}^{(E)}$ | $d_E$ | 0.16 s |
| Shape of $F(I_n^{(I)})$ around $I_{thr}^{(I)}$ | $d_I$ | 0.087 s |
| Excitatory kinetic parameter | $\gamma$ | 0.641 |
| Amplitude of uncorrelated Gaussian noise $v_n$ | $\sigma$ | 0.01 nA |
| Time constant of NMDA | $\tau_{NMDA}$ | 100 ms |
| Time constant of GABA | $\tau_{GABA}$ | 10 ms |

In all the simulations reported in this article, the above stochastic differential equations were discretized and solved using the Euler-Maruyama integration method,^55^ and using the parameter values shown in Table 1. The first 10 seconds of simulation were discarded to ensure stability of the dynamical system.

### A first-order FIC approximation

Here we provide an analytical derivation of how 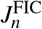 depends on other model parameters. The derivation is based on two assumptions, whose range of validity is also discussed.

As a first step in the derivation, we write down the discrete-time form of 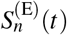 using Eq. (2e) according to the Euler-Maruyama method, which gives

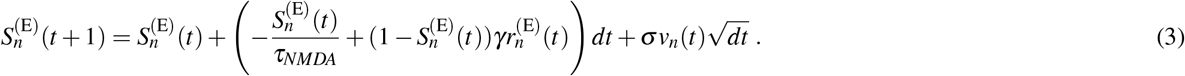

Then, we calculate the steady-state average of 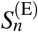 by computing the expected value to Eq. (3). Using the steady-state property, and introducing the shorthand notation 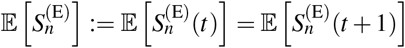, a direct calculation shows that:

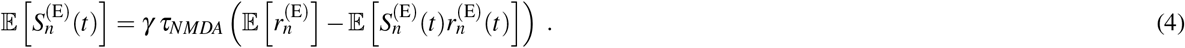

To proceed further in the derivation one needs an expression for 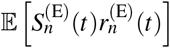. It can be found via numerical simulations that the covariance between these two variables is lower than 0.1, so we assume that 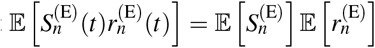 (**assumption 1**). Plugging this relationship into Eq. (4) yields

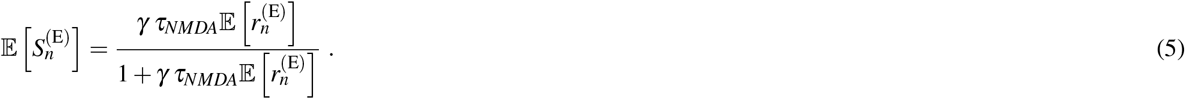

The FIC is assumed to successfully regulate the average excitatory firing rate if, for any *G*, they remain close to the firing rates obtained for the disconnected case 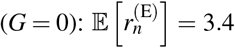 Hz for all the nodes of the network. Importantly, this implies that 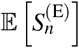 is also constant across nodes, and using Eq. (5) one finds that 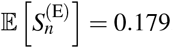. Numerical simulations for uncoupled regions give a steady-state average of 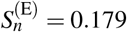, providing empirical confirmation of the accuracy of this approximation for that scenario.

As a second step of the derivation, let’s study the expected values of 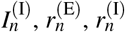, and 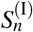 using the equations provided in the previous section. The equations for 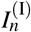 and 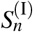 are linear, so computing the expected value is straightforward; in contrast, 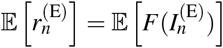 involves a non-linear function *F*. To move forward, we adopt a first-order Taylor expansion and assume that the approximation 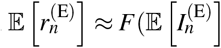 is accurate (**assumption 2**). By doing this, one obtains a system of four non-linear equations with four unknowns (the expected values of 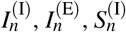 and 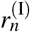), which can be solved numerically. Since the inputs to this system of equations — i.e. 𝔼 [*r*^(E)^] and 𝔼 [*S*^(E)^] — adopt the same values for all regions, the results yield the same values for all regions.

As a final step, we apply the expected value of Eq. (2a). Using the fact that 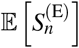 does not depend on *n*, a direct calculation shows that

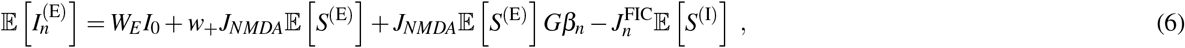

where *β*_*n*_ := ∑_*p*_ *C*_*np*_ is the strength of node *n*. To conclude, one can solve the above equation for 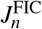 to obtain

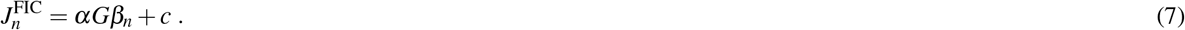

Above, 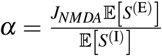 is a ratio between the expected fraction of open NMDA to GABA channels — which represents a global E/I balance parameter, and 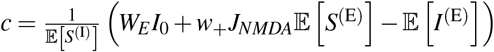 is an offset parameter that corresponds to the 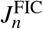 for uncoupled regions. To find values of *α* and *c*, one can plug 𝔼 [*r*^(E)^] = 3.4 Hz, the aforementioned expected values obtained for uncoupled regions, and the usual model parameters, to obtain *α* = 0.67 and *c* = 0.97. To match the value of 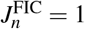 for uncoupled regions (corresponding to *G* = 0), one can use approximately *c* = 1.

Please note that this first-order approximation is based on the expected values obtained for uncoupled regions, so one should expect that **assumption 2** may become less accurate as *G* increases, which reflects that the non-linear effects of *F*(·) play a more important role in determining the optimal value of 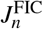 to attain stability.

### Resting-State fMRI signals

#### Participants

A total of 63 healthy subjects (36 females, mean ± SD, 23 ± 43.3 years) were selected from a data set previously described in a sleep-related study by Tagliazucchi and Laufs^56^. Participants entered the scanner at 7 PM and were asked to relax, close their eyes, and not fight the sleep onset. We selected 13 subjects with contiguous resting state time series of at least 200 volumes to perform our analysis. The local ethics committee approves the experimental protocol (Goethe-Universität Frankfurt, Germany, protocol number: 305/07), and written informed consent was asked to all participants before the experiment. The study was conducted according to the Helsinki Declaration on ethical research.

#### MNI data acquisition

MRI images were acquired on a 3-T Siemens Trio scanner (Erlangen, Germany) and fMRI acquisition parameters were 1505 volumes of T2-weighted echo planar images, *TR/TE* = 2080 *ms/*30 *ms*, matrix 64 × 64, voxel size 3 × 3 × 3 *mm*^3^, distance factor 50%; FOV 192 *mm*^2^.

#### Brain parcellation to extract BOLD time series

##### AAL 90

To extract the time series of BOLD signals from each participant in a coarse parcellation, we used the AAL90 parcellation with 90 brain areas anatomically defined in Tzourio-Mazoyer et al., 2002.^36^

##### Schaefer 1000

To extract the time series of BOLD signals from each participant, in a finer scale we used the Schaefer functional parcellation with 1000 brain areas, which was based on estimation from a large dataset (n = 1489).^47^

##### Filtering

BOLD signals (empirical or simulated) were filtered with a Butterworth (order 2) band-pass filter in the 0.01-0.1 frequency range.

### Functional Connectivity and Functional Connectivity Dynamics

The Functional Connectivity (FC) matrix was obtained computing the Pearson correlation coefficient between all the pairs of simulated or recorded BOLD signals. The Functional Connectivity Dynamics (FCD) was obtained by computing the FC(t), where *t* is given by consecutive sliding windows of length TR=30 and TR=28 of overlap. Then, we vectorized each of the FC(t) by taking the upper triangular, and finally computed the Pearson correlation coefficient between all these vectors to obtain the FCD matrix.

### Bayesian Optimization

A Matlab implementation of Bayesian Optimization^43,44^ with an expected-improvement acquisition function was used to optimize the DMF model parameters. The objective function was defined as the Kolmogorov-Smirnov (K-S) distance between the histogram of the pooled empirical Functional Connectivity Dynamics (*FCD*_*emp*_) and the histogram of the simulated FCD (*FCD*_*dmf*_). We simulated 500 seconds of BOLD signals sampled at TR=2, generating a number of time points comparable to the empirical BOLD signals. The optimization was run assuming a stochastic objective function, letting the algorithm to randomly select the initial conditions for each simulation.

### Structural Connectivity

#### AAL 90

The structural connectome was obtained applying diffusion tensor imaging (DTI) to diffusion weighted imaging (DWI) recordings from 16 healthy right-handed participants (11 men and 5 women, mean age: 24.75 *±* 2.54 years) recruited online at Aarhus University, Denmark, as used in previous studies.^12,16,57^ Briefly, the construction of the structural connectivity matrix (SC) was performed following the following steps: the regions defined using the AAL template^36^ were warped from MNI space to the diffusion MRI native space using the FLIRT tool from the FSL toolbox (www.fmrib.ox.ac.uk/fsl, FMRIB, Oxford). Then, probabilistic tractography with default parameters of the FSL diffusion toolbox (Fdt) were used to estimate the connections between region. The local probability distribution of fibre direction at each voxel was estimated following Behrens et al. (2003)^58^ and the probtrackx tool in Fdt was used for the automatic estimation of crossing fibres within each voxel. Using a sampling of 5000 streamlines per voxel, the connection probability from a seed voxel *i* to another voxel j was defined as the proportion of fibres flowing through voxel *i* that reach voxel *j*.^59^ The fraction of sampled fibres in all voxels in a region *i* that reach any voxel in region *j* in an AAL region *i* determines the connection probability *P*_*i j*_ between those regions. Due to the dependence of tractography on the seeding location, the probability *P*_*i j*_ is not necessarily equivalent to *P*_*ji*_. However, these two probabilities are highly correlated across the brain for all participants (the least Pearson *r* = 0.70, p *<* 10 50), and thus the unidirectional connectivity probability *P*_*i j*_ between regions i and j was defined by averaging these two connectivity probabilities. This connectivity was considered as a measure of the structural connectivity resulting in a 90×90 symmetric weighted matrix C representing the network organization of each individual brain. A group averaged structural connectivity matrix was obtained by averaging across all 16 healthy participants.

#### Schaefer 1000

As described in Ref.^1^, we used the Human Connectome Project (HCP) database, which contains diffusion spectrum and T2-weighted neuroimaging data. Specifically, we estimated the structural connectivity using HCP dMRI dataset provided by the Special HCP dMRI, which uses excellent protocols taking 89 minutes for each of 32 HCP participants at the MGH centre. A detailed description of the acquisition parameters can be found in the HCP website.^60^. The precise preprocessing is described in details in Horn and colleagues^61^, In brief, the data were processed by using a q-sampling imaging algorithm implemented in DSI studio (http://dsi-studio.labsolver.org). A white-matter mask was computed by segmenting the T2-weighted images and co-registering the images to the b0 image of the diffusion data using SPM12. 200.000 fibres were sampled within the white-matter mask for each HCP participant. Fibres were transformed into MNI space using Lead-DBS.^62^ Finally, we used the standardized methods in Lead-DBS to extract the structural connectomes from the Schaefer-1000 parcellation.^47^ A group averaged structural connectivity matrix was obtained by averaging across all 32 healthy participants.

## Data and code availability

The C++ implementation of the FastDMF, its Python and Matlab wrappers, and the corresponding data sets can be found at https://gitlab.com/concog/fastdmf

## Acknowledgements

R.H. is funded by CONICYT scholarship CONICYT-PFCHA/Doctorado Nacional/2018-21180428. P.M. is funded by the Wellcome Trust (grant no. 210920/Z/18/Z). F.R. is supported by the Ad Astra Chandaria Foundation. A.I.L. is funded by a Gates Cambridge Scholarship (OPP 1144). E.T is funded by Agencia Nacional De Promocion Cientifica Y Tecnologica (Argentina) (Grant No. PICT-2018-03103). R.C. was supported by the Human Brain Project, H2020-945539. G.D. is supported by the Spanish Research Project (ref. PID2019-105772GB-I00 AEI FEDER EU), funded by the Spanish Ministry of Science, Innovation and Universities (MCIU), State Research Agency (AEI) and European Regional Development Funds (FEDER); HBP SGA3 Human Brain Project Specific Grant Agreement 3 (Grant Agreement No. 945539), funded by the EU H2020 FET Flagship program and SGR Research Support Group support (ref. 2017 SGR 1545), funded by the Catalan Agency for Management of University and Research Grants (AGAUR).

## Author contributions statement

R.H. performed the numerical analysis and run the simulations. All the authors of this article conceptualized this research, design the methodology. All the authors carefully analysed the results, drafted and reviewed the manuscript.

## Footnotes

1 In other words, the DMF equations are used to simulate firing rates, which are temporarily stored in a buffer, and then consumed by the BK integrator to generate the slow BOLD signals. Once a batch of firing rates has been consumed by the BK integrator, the DMF integrator is notified and allowed to write on the same memory address, and the cycle begins again. This mechanism makes the total memory usage of the simulation grow with the size of the resulting BOLD signals, as opposed to the firing rate signals, which are approximately 1000 times larger in memory.

2 We divide it by 2 to compensate the double counting of the symmetric entries of *C*

## References

1. Deco, G. & Kringelbach, M. L. Turbulent-like dynamics in the human brain. Cell Reports 33, 108471, DOI: 10.1016/J.CELREP.2020.108471 (2020).

2. Munn, B. R., Müller, E. J., Wainstein, G. & Shine, J. M. The ascending arousal system shapes neural dynamics to mediate awareness of cognitive states. Nat. Commun. 12, DOI: 10.1038/s41467-021-26268-x (2021).

3. Cofré, R. et al. Whole-brain models to explore altered states of consciousness from the bottom up. Brain Sci. 10, DOI: 10.3390/brainsci10090626 (2020).

4. Fox, K. C. et al. Intrinsic network architecture predicts the effects elicited by intracranial electrical stimulation of the human brain. Nat. Hum. Behav. 4, DOI: 10.1038/s41562-020-0910-1 (2020).

5. B., S. W. & Milner, B. Loss of recent memory after bilateral hippocampal lesions. J. neurology, neurosurgery, psychiatry 20, DOI: 10.1136/jnnp.20.1.11 (1957).

6. Damasio, H., Grabowski, T., Frank, R., Galaburda, A. M. & Damasio, A. R. The return of Phineas Gage: Clues about the brain from the skull of a famous patient. Science 264, DOI: 10.1126/science.8178168 (1994).

7. Bisiach, E. & Luzzatti, C. Unilateral Neglect of Representational Space. Cortex 14, DOI: 10.1016/S0010-9452(78)80016-1 (1978).

8. Weiskrantz, L. Outlooks for blindsight: Explicit methodologies for implicit processes, DOI: 10.1098/rspb.1990.0016 (1990).

9. Miesenböck, G. Optogenetic control of cells and circuits. Annu. Rev. Cell Dev. Biol. 27, DOI: 10.1146/annurev-cellbio-100109-104051 (2011).

10. Deco, G., Hagmann, P., Romani, G. L., Mantini, D. & Corbetta, M. How local excitation-inhibition ratio impacts the whole brain dynamics. J. Neurosci. 34, 7886–7898, DOI: 10.1523/JNEUROSCI.5068-13.2014 (2014).

11. Cabral, J., Kringelbach, M. L. & Deco, G. Functional connectivity dynamically evolves on multiple time-scales over a static structural connectome: Models and mechanisms, DOI: 10.1016/j.neuroimage.2017.03.045 (2017).

12. Deco, G. et al. Whole-brain multimodal neuroimaging model using serotonin receptor maps explains non-linear functional effects of LSD. Curr. Biol. 1–10, DOI: 10.1016/j.cub.2018.07.083 (2018).

13. Deco, G. et al. Perturbation of whole-brain dynamics in silico reveals mechanistic differences between brain states. NeuroImage DOI: 10.1016/j.neuroimage.2017.12.009 (2018).

14. Deco, G. et al. Awakening: Predicting external stimulation to force transitions between different brain states. Proc. Natl. Acad. Sci. United States Am. DOI: 10.1073/pnas.1905534116 (2019).

15. Herzog, R. et al. A mechanistic model of the neural entropy increase elicited by psychedelic drugs. Sci. Reports 1–12, DOI: 10.1101/2020.05.13.093732 (2020).

16. Ipiña, I. P. et al. Modeling regional changes in dynamic stability during sleep and wakefulness. NeuroImage DOI: 10.1016/j.neuroimage.2020.116833 (2020).

17. Kringelbach, M. L. et al. Dynamic coupling of whole-brain neuronal and neurotransmitter systems. Proc. Natl. Acad. Sci. United States Am. 117, 9566–9576, DOI: 10.1073/pnas.1921475117 (2020).

18. Deco, G., Tononi, G., Boly, M. & Kringelbach, M. L. Rethinking segregation and integration: Contributions of whole-brain modelling, DOI: 10.1038/nrn3963 (2015).

19. Goldman, J. S. et al. Bridging Single Neuron Dynamics to Global Brain States. Front. Syst. Neurosci. DOI: 10.3389/fnsys.2019.00075 (2019).

20. Shine, J. M. et al. Computational models link cellular mechanisms of neuromodulation to large-scale neural dynamics, DOI: 10.1038/s41593-021-00824-6 (2021).

21. Coronel-Oliveros, C., Cofré, R. & Orio, P. Cholinergic neuromodulation of inhibitory interneurons facilitates functional integration in whole-brain models. PLoS Comput. Biol. DOI: 10.1371/JOURNAL.PCBI.1008737 (2021).

22. Breakspear, M. Dynamic models of large-scale brain activity, DOI: 10.1038/nn.4497 (2017).

23. Deco, G. & Kringelbach, M. L. Great expectations: Using whole-brain computational connectomics for understanding neuropsychiatric disorders, DOI: 10.1016/j.neuron.2014.08.034 (2014).

24. Ritter, P., Schirner, M., Mcintosh, A. R. & Jirsa, V. K. The Virtual Brain Integrates Computational Modeling and Multimodal Neuroimaging. Brain Connect. DOI: 10.1089/brain.2012.0120 (2013).

25. Cakan, C., Jajcay, N. & Obermayer, K. neurolib: A Simulation Framework for Whole-Brain Neural Mass Modeling. Cogn. Comput. DOI: 10.1007/s12559-021-09931-9 (2021).

26. Burt, J. B. et al. Transcriptomics-informed large-scale cortical model captures topography of pharmacological neuroimaging effects of lsd. eLife 10, DOI: 10.7554/eLife.69320 (2021).

27. Deco, G. et al. Resting-state functional connectivity emerges from structurally and dynamically shaped slow linear fluctuations. J. Neurosci. DOI: 10.1523/JNEUROSCI.1091-13.2013 (2013).

28. Van Vreeswijk, C. & Sompolinsky, H. Chaos in neuronal networks with balanced excitatory and inhibitory activity. Science DOI: 10.1126/science.274.5293.1724 (1996).

29. Amit, D. J. & Brunel, N. Model of global spontaneous activity and local structured activity during delay periods in the cerebral cortex. Cereb. Cortex DOI: 10.1093/cercor/7.3.237 (1997).

30. Stephan, K. E., Weiskopf, N., Drysdale, P. M., Robinson, P. A. & Friston, K. J. Comparing hemodynamic models with DCM. NeuroImage 38, 387–401, DOI: 10.1016/j.neuroimage.2007.07.040 (2007).

31. Demirtaş, M. et al. Hierarchical Heterogeneity across Human Cortex Shapes Large-Scale Neural Dynamics. Neuron DOI: 10.1016/j.neuron.2019.01.017 (2019).

32. Wang, P. et al. Inversion of a large-scale circuit model reveals a cortical hierarchy in the dynamic resting human brain. Trop. Subtrop. Agroecosystems DOI: 10.1126/sciadv.aat7854 (2019).

33. Luppi, A. I. et al. Paths to oblivion: Common neural mechanisms of anaesthesia and disorders of consciousness. Prepr. bioRxiv 2021.02.14.431140 (2020).

34. Gatica, M. et al. High-order functional interactions in ageing explained via alterations in the connectome in a whole-brain model. Bioarxiv DOI: 10.1101/2021.09.15.460435 (2021).

35. Hellyer, P. J., Jachs, B., Clopath, C. & Leech, R. Local inhibitory plasticity tunes macroscopic brain dynamics and allows the emergence of functional brain networks. NeuroImage DOI: 10.1016/j.neuroimage.2015.08.069 (2016).

36. Tzourio-Mazoyer, N. et al. Automated anatomical labeling of activations in SPM using a macroscopic anatomical parcellation of the MNI MRI single-subject brain. NeuroImage 15, 273–89, DOI: 10.1006/nimg.2001.0978 (2002).

37. Desikan, R. S. et al. An automated labeling system for subdividing the human cerebral cortex on mri scans into gyral based regions of interest. NeuroImage 31, 968–980, DOI: https://doi.org/10.1016/j.neuroimage.2006.01.021 (2006).

38. Deco, G. & Kringelbach, M. L. Turbulent-like Dynamics in the Human Brain. Cell Reports DOI: 10.1016/j.celrep.2020.108471 (2020).

39. Kobeleva, X., López-González, A., Kringelbach, M. L. & Deco, G. Revealing the relevant spatiotemporal scale underlying whole-brain dynamics. bioRxiv DOI: 10.1101/2020.09.12.277699 (2021).https://www.biorxiv.org/content/early/2021/03/06/2020.09.12.277699.full.pdf.

40. Guennebaud, G., Jacob, B. et al. Eigen v3. http://eigen.tuxfamily.org (2010).

41. MathWorks, T. MATLAB C++ Math Library User’s Guide, vol. User’s Guide Version 2 (The MathWorks, 1999).

42. Abrahams, D. & Grosse-Kunstleve, R.W. Building hybrid systems with Boost.Python. C/C++ Users J. 21 (2003).

43. Shahriari, B., Swersky, K., Wang, Z., Adams, R. P. & De Freitas, N. Taking the human out of the loop: A review of Bayesian optimization, DOI: 10.1109/JPROC.2015.2494218 (2016).

44. Ulmasov, D., Baroukh, C., Chachuat, B., Deisenroth, M. P. & Misener, R. Bayesian Optimization with Dimension Scheduling: Application to Biological Systems. Comput. Aided Chem. Eng. DOI: 10.1016/B978-0-444-63428-3.50180-6 (2016).

45. Hansen, E. C., Battaglia, D., Spiegler, A., Deco, G. & Jirsa, V. K. Functional connectivity dynamics: Modeling the switching behavior of the resting state. NeuroImage 105, DOI: 10.1016/j.neuroimage.2014.11.001 (2015).

46. Van Essen, D. C. et al. The wu-minn human connectome project: an overview. Neuroimage 80, 62–79 (2013).

47. Schaefer, A. et al. Local-Global Parcellation of the Human Cerebral Cortex from Intrinsic Functional Connectivity MRI. Cereb. Cortex DOI: 10.1093/cercor/bhx179 (2018).

48. Atasoy, S., Donnelly, I. & Pearson, J. Human brain networks function in connectome-specific harmonic waves. Nat. Commun. 7, 10340, DOI: 10.1038/ncomms10340 (2016).

49. Deco, G. et al. Rare long-range cortical connections enhance human information processing. Curr. Biol. DOI: 10.1016/j.cub.2021.07.064 (2021).

50. Escrichs, A. et al. Unifying turbulent dynamics framework distinguishes different brain states. bioRxiv (2021).

51. De Filippi, E. et al. The menstrual cycle modulates whole-brain turbulent dynamics. Front. Neurosci. 15, 1658, DOI: 10.3389/fnins.2021.753820 (2021).

52. Naskar, A. et al. Multi-scale dynamic mean field model (MDMF) relates resting-state brain dynamics with local cortical excitatory-inhibitory neurotransmitter homeostasis. Netw. Neurosci. DOI: https://doi.org/10.1162/netn_a_00197 (2021).

53. Kong, X. et al. Sensory-motor cortices shape functional connectivity dynamics in the human brain. Nat. Commun. DOI: 10.1038/s41467-021-26704-y (2021).

54. Wong, K. F. & Wang, X. J. A recurrent network mechanism of time integration in perceptual decisions. J. Neurosci. DOI: 10.1523/JNEUROSCI.3733-05.2006 (2006).

55. Kloeden, P. E. & Platen, E. Numerical Solution of Stochastic Differential Equations, vol. 23 (Springer, 2013).

56. Tagliazucchi, E., Carhart-Harris, R., Leech, R., Nutt, D. & Chialvo, D. R. Enhanced repertoire of brain dynamical states during the psychedelic experience. Hum. Brain Mapp. 35, 5442–5456, DOI: 10.1002/hbm.22562 (2014).

57. Cabral, J., Hugues, E., Kringelbach, M. L. & Deco, G. Modeling the outcome of structural disconnection on resting-state functional connectivity. NeuroImage 62, 1342–1353, DOI: 10.1016/J.NEUROIMAGE.2012.06.007 (2012).

58. Behrens, T. E. et al. Characterization and propagation of uncertainty in diffusion-weighted mr imaging. Magn. Reson. Medicine: An Off. J. Int. Soc. for Magn. Reson. Medicine 50, 1077–1088 (2003).

59. Behrens, T. E., Berg, H. J., Jbabdi, S., Rushworth, M. F. & Woolrich, M. W. Probabilistic diffusion tractography with multiple fibre orientations: What can we gain? NeuroImage 34, 144–155, DOI: 10.1016/j.neuroimage.2006.09.018 (2007).

60. Setsompop, K. et al. Pushing the limits of in vivo diffusion mri for the human connectome project. NeuroImage 80, 220–233 (2013).

61. Horn, A., Neumann, W.-J., Degen, K., Schneider, G.-H. & Kühn, A. A. Toward an electrophysiological “sweet spot” for deep brain stimulation in the subthalamic nucleus. Hum. brain mapping 38, 3377–3390 (2017).

62. Horn, A. & Blankenburg, F. Toward a standardized structural–functional group connectome in mni space. NeuroImage 124, 310–322, DOI: 10.1016/J.NEUROIMAGE.2015.08.048 (2016).

